# Random forest model improves annotation and discovery of variants of uncertain significance in Alzheimer’s and other neurological disorders

**DOI:** 10.1101/2025.10.02.680068

**Authors:** Caroline Jonson, Mary B. Makarious, Mathew J. Koretsky, Dan Vitale, Liya Rabkina, Argentina Lario Lago, Manizhe Eslami-Amirabadi, Eliana Marisa Ramos, Priyanka Narayan, Jennifer S. Yokoyama, Andrew B. Singleton, Cornelis Blauwendraat, Mark R. Cookson, Mike A. Nalls, Hampton L. Leonard

**Affiliations:** Center for Alzheimer’s and Related Dementias, Bethesda, MD, USA 20892; DataTecnica LLC, Washington, DC, USA 20037; Laboratory of Neurogenetics, NIA, NIH, Bethesda, MD, USA 20892; Memory and Aging Center, Department of Neurology, Weill Institute for Neurosciences, University of California, San Francisco, CA, USA; Department of Radiology and Biomedical Imaging, University of California, San Francisco, CA, USA; University of Washington, School of Medicine, Memory and Brain Wellness Center at Harborview, Seattle, WA, 98104; Department of Neurology, David Geffen School of Medicine, University of California, Los Angeles, Los Angeles, CA, USA; Coalition for Aligning Science, Chevy Chase, MD, USA; Global Parkinson’s Genetics Program, Chevy Chase, MD, USA; Aligning Science Across Parkinson’s (ASAP), Chevy Chase, MD, USA

**Author notes:** denotes corresponding authorship **Materials and correspondence** Correspondence should be directed to Mike A. Nalls and Hampton L. Leonard. denotes equal contribution (Caroline Jonson, Mary B. Makarious).

## Abstract

Variants of uncertain significance (VUS) are a bottleneck for genetic discovery and complicate clinical decision-making in Alzheimer’s disease and related neurological disorders (ADRD). We developed MoVUS: Model for Variants of Unknown Significance, a random-forest approach that integrates functional predictors to classify missense VUS.

MoVUS leverages a balanced random forest model trained on dbNSFP v5.1a with high-confidence ClinVar and HGMD labels, using harmonized functional prediction rankscores. MoVUS produced confident, explainable calls, with ∼98% accuracy (AUC ∼0.998), prioritizing potentially pathogenic candidates and down-ranked likely benign variants on independent validation sets of ClinVar-only and HGMD-only variants. In our discovery analyses on ADRD-implicated variants in dbNSFP and from independent collaborator cohorts, we achieved high-confidence classifications on a majority of the unknown variants (average of 55% of discovery variants). We also had access to medical records and family trees for some variants, further validating our findings.

Across held-out and external datasets, MoVUS reports high accuracy alongside confidence scores and helps prioritize actionable candidates, and reduces bias by considering multiple scores for each variant. To facilitate use, we developed a web app for users to browse across 100+ ADRD genes. MoVUS provides transparent, reproducible triage for research follow-up by pairing consensus predictors with SHAP-based visualizations and explanations.

## Introduction

Variants of unknown significance (VUS) present a significant challenge in genetic interpretation, as their clinical relevance often remains unresolved. Traditional predictive models frequently struggle to accurately classify these variants due to the complexity of genomic data and the many factors influencing pathogenicity. Though numerous annotation resources have been developed to predict the functional impact and pathogenicity of variants, integrating these diverse data sources in a meaningful way remains a significant obstacle for researchers and clinicians.

To address this challenge, we developed MoVUS (Model for Variants of Unknown Significance), a machine learning pipeline designed to classify VUS by leveraging large-scale genomic annotation resources. MoVUS systematically integrates 25 harmonized functional prediction scores (summarized in **Table S2**) from dbNSFP v5.1a, including conservation/constraint measures (CADD^1^, GERP^2^, Eigen-PC^3^), deep learning models (AlphaMissense^4^, PHACTboost^5^, ESM-1b^6^), functional contraint tools (SIFT4G^7^, PolyPhen-2^8^, MutationTaster^9^), meta-predictors (ClinPred^10^, REVEL^11^, PrimateAI^12^), and phylogenetic conservation scores (phyloP 100-way vertebrate conservation^13^) to train robust classifiers that distinguish pathogenic from benign variants.

Different annotation tools generate scores using methodologies that capture unique and complementary aspects of variant impact. By applying a random forest model that leverages these scores, MoVUS provides a more comprehensive and accurate pathogenicity assessment, particularly in the context of disease risk.

Here, we introduce MoVUS and demonstrate its application on independent test datasets to identify potentially novel variants implicated in neurodegenerative disease. Our results highlight the potential of integrative machine learning approaches to improve variant interpretation. **Figure 1** provides a graphical overview of the MoVUS workflow, illustrating the datasets used for model training, independent testing, and the discovery of potentially pathogenic VUS.

**Figure 1.**
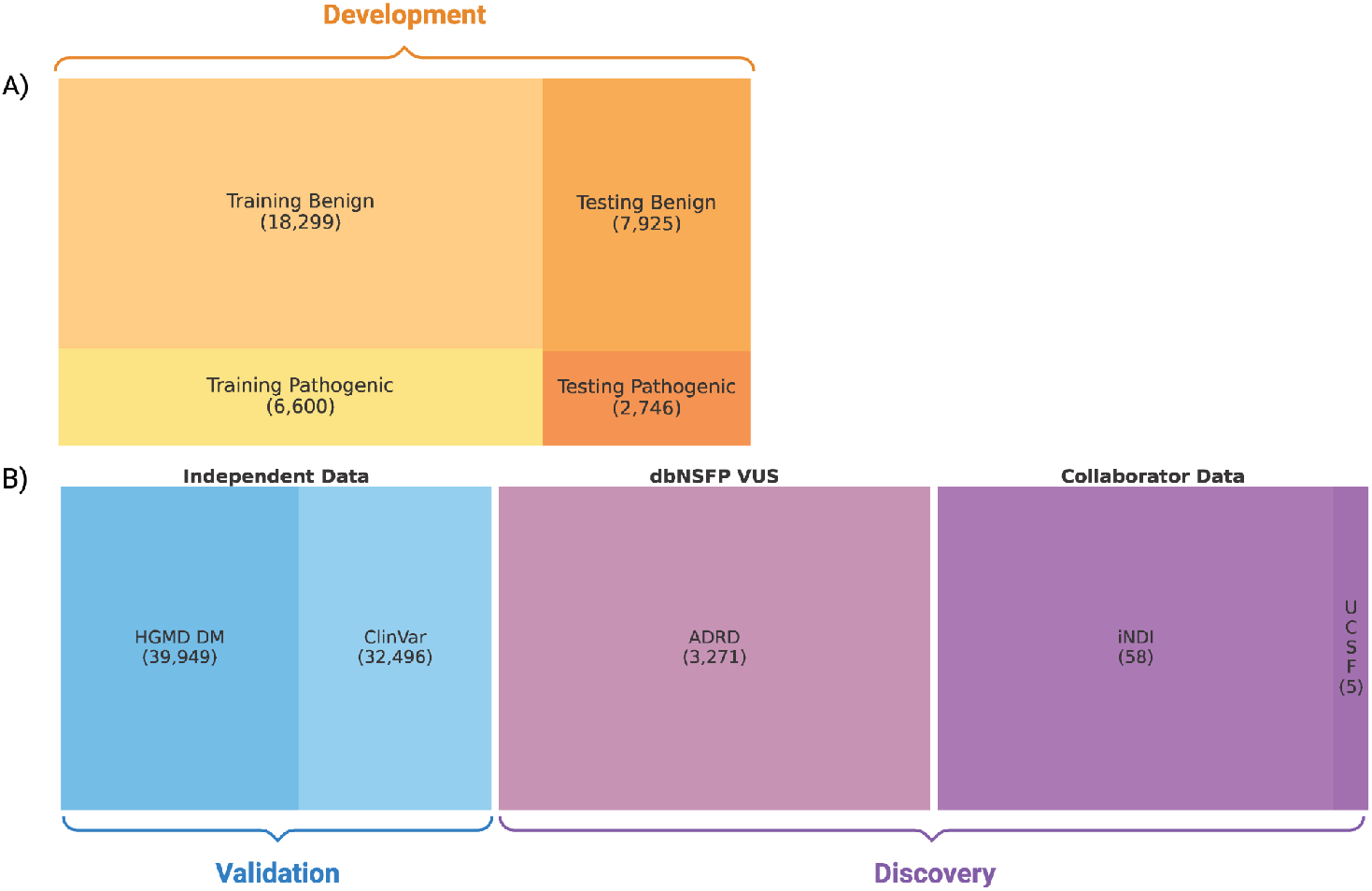
MoVUS overview of Variant Datasets for Model Development, Validation, and Discovery. A) The development dataset (dbNSFP v5.1a) consists of training and testing data with a 70:30 split. B) The validation and discovery datasets include independent validation data (HGMD DM and ClinVar) and discovery sets (dbNSFP VUS ADRD variants and Collaborator Data from INDI and UCSF). Square areas are proportional to variant counts.

## Materials and Methods

### Data Sources

#### dbNSFP acquisition and preprocessing

MoVUS was trained using data from the database for nonsynonymous SNPs’ functional predictions (dbNSFP v5.1a; https://www.dbnsfp.org). This database has compiled pathogenicity prediction scores, conservation scores, and other variant annotations from over 40 sources on a total of 83,572,434 non-synonymous (nsSNVs) and splice site variants (ssSNVs), including ClinVar classifications for pathogenicity of germline variants for Mendelian diseases. All variant coordinates in dbNSFP are mapped to the GRCh38/hg38 human reference genome assembly. The dbNSFP file was downloaded from https://sites.google.com/site/jpopgen/dbNSFP (March 2025).

For model training, we focused exclusively on variants labeled with ClinVar classifications of ‘Pathogenic’ and ‘Benign’ rather than including ‘Likely Pathogenic’ or ‘Likely Benign’ variants with at least a single submitter criterion. This conservative approach ensures high-quality training labels for both classes, minimizes label noise, and provides a more robust foundation for learning the distinct patterns of definitively pathogenic and benign variants. Variants not belonging to these categories were reclassified as Variants of Unknown Significance (VUS) and removed from downstream analyses. The most deleterious consequence per annotation was preserved for variants with duplicate rows.

We obtained additional clinical variant classifications from the Human Gene Mutation Database (HGMD, Professional version), accessed under institutional license. Collaborator-provided datasets, including variants of unknown significance (VUS) from neurodegenerative disease cohorts, were used for independent validation and discovery analyses.

### Variant Annotation and Preprocessing

dbNSFP v5.1a included all variant annotations, and data preprocessing was performed on this dataset. Sequence Ontology (SO) labels were assigned to each variant based on amino acid changes, using a custom Python script that classified variants as missense, stop gained, start lost, start gained, stop lost, or splice donor/acceptor as outlined in **Table S2**. For all downstream analyses, we retained only missense variants.

Relevant features were extracted for further analysis, including functional prediction scores, SO labels, and clinical annotations. Only rankscore columns were used for functional prediction scores, scaled between 0 and 1, and thus harmonized across all other annotations. To reduce multicollinearity, we computed the pairwise correlation matrix for all features (**Figure S1**). For any pair of features with a correlation coefficient greater than 0.9, we retained only one feature from the pair, prioritizing those with higher biological relevance or predictive performance, as assessed by our single-feature performance assessment, to avoid overfitting. This resulted in the removal of 16 features. All annotations included or the reason for their exclusion are summarized in **Table S1**.

For functional prediction rankscores, any remaining missing values were imputed using the median value for each feature. All selected rankscore features were retained for model development since none exceeded the pre-specified missingness threshold (>50%). In contrast, all allele frequency columns were removed due to their extremely high levels of missingness.

### Dataset Construction and Splitting

HGMD data were merged with dbNSFP by matching on chromosome, position, reference, and alternate alleles. For variants present in both datasets, HGMD classification scores (e.g., “DM” for disease-causing mutation, “DP” for disease-associated polymorphism, “FP” for functional polymorphism, and “DFP” for disease-associated polymorphism) were added to the dbNSFP annotations. Only variants classified as DM in HGMD were retained for model development and evaluation to be consistent with only including “Pathogenic” or “Benign” classified variants from Clinvar. Variants in only one dataset (i.e., not shared between HGMD and dbNSFP) were retained for independent testing and discovery analyses.

The merged dataset was randomly split into training and test sets using a 70:30 ratio without stratification because we didn’t need to force equal distribution of target classes in each set. Additional independent test sets were constructed from variants unique to HGMD or ClinVar, and discovery analyses were performed on external VUS datasets from collaborators.

### VUS datasets from collaborators

#### UCSF VUS

The five variants provided by the University of California, San Francisco (UCSF) were manually curated to highlight the most interesting novel variants of uncertain significance (VUS) in major Alzheimer’s disease (AD) and frontotemporal dementia (FTD) genes. These variants were prioritized based on predicted deleteriousness by in silico tools, location within mutation hotspots, and/or prior reports in the literature with inconclusive evidence. They were drawn from a dataset of variants identified over several years through UCSF targeted gene panels and/or whole-exome sequencing collected from patients assessed and clinically diagnosed at the University of California, San Francisco Memory and Aging Center (UCSF MAC)^14,15^. None of the participants in these studies carried any known disease-causing pathogenic variant. The cohorts from which these variants were discovered, include individuals with diverse forms of FTD and related conditions, such as corticobasal syndrome (CBS), progressive supranuclear palsy (PSP), amyotrophic lateral sclerosis (ALS), AD, behavioral variant FTD (bvFTD), primary progressive aphasia (PPA), corticobasal degeneration (CBD), Creutzfeldt–Jakob disease (CJD), dementia with Lewy bodies (DLB), chronic traumatic encephalopathy (CTE), mild cognitive impairment (MCI), and other neurological disorders affecting cognition, excluding vascular dementia.

#### iNDI VUS

The 58 variants that were used to develop the IPSC neurodegenerative disease initiative (iNDI) from the Center for Alzheimer’s and Related Dementias (CARD) were initially taken from the Human Gene Mutation Database, focusing on genes with variants where dementia has been reported in the human phenotype, selecting multiple variants where possible, and prioritizing variants with the highest numbers of publications. These variants remained from 229 original iNDI variants after filtering the variants already listed in our dbNSFP ADRD-related VUS discovery set to ensure all datasets were independent and not overlapping.

### Model Development

Model training and evaluation leveraged scikit-learn (v1.6.1) and imbalanced-learn (v0.13.0). More specifically, Hyperparameter optimization was achieved using 5-fold cross-validation with GridSearchCV, optimizing for balanced accuracy. For variant classification, we used a BalancedRandomForestClassifier from the imbalanced-learn library ^16^. The final model was trained on the complete training set after CV using the optimal hyperparameters identified through grid search (**Table S3**).

### Model Evaluation and Discovery

Independent validation was performed on external datasets comprising HGMD-only and ClinVar-only variants separately. Conservative probability thresholds of 0.35 and 0.65 were selected to maximize data inclusion while maintaining model performance superior to the existing alternative scores, reporting scores between these probabilities as inconclusive to the end-user for that variant. The model was also applied to discovery datasets, including ADRD-related VUS from ClinVar and collaborator VUS datasets (UCSF and iNDI, described above) to assess performance on variants of unknown significance with more clinically relevant high-confidence prediction thresholds of 0.1 and 0.9, with variants falling between these thresholds again classified as uncertain. Stricter thresholds were used here, as minimizing incorrectly classified variants is a larger priority for the discovery set than maximizing the amount of variants classified. Variants that overlapped with those used in model training, other test sets, or internal discovery sets were removed from each collaborator dataset to ensure an independent evaluation.

Individual feature analysis served as a benchmark to assess each annotation’s predictive performance independently and demonstrate that MoVUS outperforms any single annotation. For each feature, a RandomForestClassifier was trained using only that single feature, with 5-fold cross-validation on the training set and evaluation on the test set.

Each test set’s performance metrics were calculated separately to assess model robustness across different data sources and variant classifications.

### Model Explainability

SHAP (SHapley Additive exPlanations) was used for model interpretability, providing global feature importance rankings and local explanations for individual predictions ^17^. In other words, SHAP analysis shows us the overall predictive importance of each feature during training and how each feature contributes to the pathogenicity prediction for any given variant. SHAP plots for each variant are included in our Streamlit app (referenced below).

### Data and Code Availability

All code, including environment files with exact package versions, is available on GitHub under the Apache 2.0 license [https://github.com/NIH-CARD/MoVUS]. To maximize accessibility, we developed a web-based interface using Streamlit and deployed it on Google Cloud Run. The implementation focuses on 102 ADRD-related genes, providing interactive threshold adjustment (Model Default: 0.49/0.51, Clinical: 0.1/0.9, Conservative: 0.35/0.65) and comprehensive variant search functionality. The application incorporates SHAP visualization for model interpretability and implements intelligent result limiting (500 variants maximum) for optimal performance. The web interface is publicly accessible at https://movus-variant-explorer-441353018952.us-central1.run.app and serves as a practical demonstration of MoVUS’s utility in real-world research scenarios.

## Results

### Model Training and Evaluation

MoVUS was trained using data from the database for nonsynonymous SNPs’ functional predictions (dbNSFP). A total of 35,570 missense variants with high-confidence classifications were included in the analysis. The dataset was randomly split into training (70%) and test (30%) sets. The training set comprised 24,899 variants, including 18,299 benign and 6,600 pathogenic variants. The test set included 10,671 variants, of which 7,925 were benign and 2,746 were pathogenic. This split was used for all model development and evaluation.

The model achieved a 5-fold cross-validation balanced accuracy of 98.4% ± 0.002 on the training set. MoVUS achieved a balanced accuracy of 98.4% on the held-out test set with an AUC of 99.8%. Predictions (probability thresholds of 0.35 and 0.65) were achieved for 10,492 out of 10,671 test variants (98.3%), with a balanced accuracy of 98.9% for these predictions. The remaining variants between these thresholds remained unclassified. Detailed performance metrics are provided in **Table 1**.

**Table 1.**
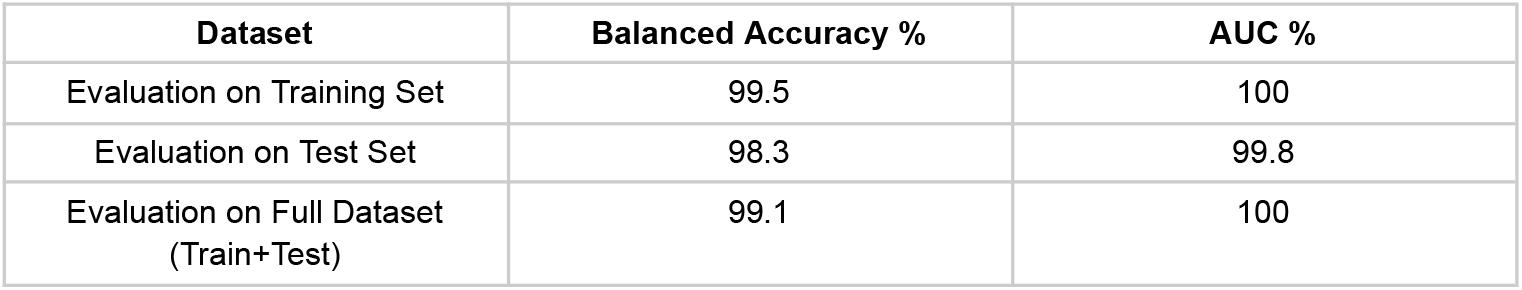
MoVUS development metrics following training and testing.

### Feature Importance and Model Interpretation

SHAP analysis revealed that MoVUS relies most heavily on consensus-based prediction tools, as the top 4 contributing features are ensemble methods that integrate multiple individual prediction algorithms: MutationTaster, ClinPred, MutScore, and PHACTboost (**Figure 2**). This suggests that MoVUS successfully recognizes and leverages the value of pre-validated consensus approaches, creating a meta-consensus score that combines the best available evidence from multiple prediction algorithms.

**Figure 2.**
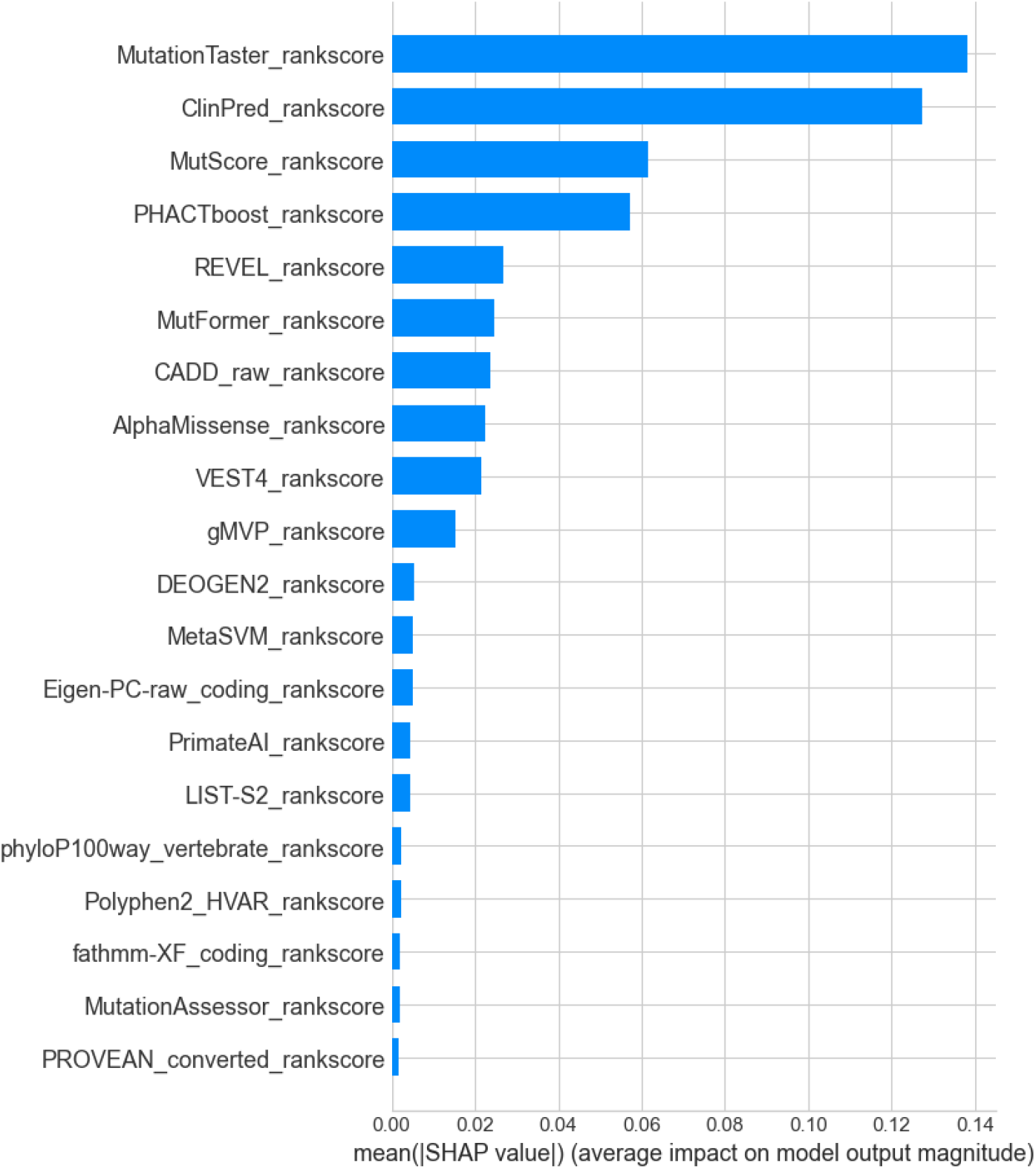
SHAP Feature Importance for MoVUS. This barplot highlights the feature importance ranking, showing the average impact of computational prediction tools on the model’s output magnitude, measured by mean absolute SHAP value, and is sorted from most important to least important.

To validate our feature selection and understand the contribution of each annotation independently, we evaluated the performance of each feature in isolation. For each of the 25 features in MoVUS, we trained a Random Forest classifier using only that single feature, with 5-fold cross-validation on the training set and evaluation on the independent ClinVar test set.

Unsurprisingly, individual feature analysis also revealed that consensus-based prediction tools outperformed individual functional prediction scores, with four of the top five performing features being ensemble methods that integrate multiple prediction algorithms (Table S4). ClinPred achieved the highest performance with a balanced accuracy of 94.5%, followed by MutScore, PHACTboost, and MutationTaster. The dominance of consensus tools in the top performers further validates MoVUS’s ensemble methodology of combining multiple prediction algorithms.

### Performance on Independent Test Sets

MoVUS was further evaluated on independent test sets derived from HGMD-only and ClinVar-only variants. The model maintained robust performance (with lower and upper prediction thresholds of 0.35 and 0.65, respectively to maximize both data inclusion and balanced accuracy) across these external datasets, with 92.2% recall for HGMD-only (with 7% of variants falling outside the prediction thresholds) and a balanced accuracy of 98.6% (with 98% of variants included) for ClinVar-only test sets. Predictions were achieved for the majority of variants in each set, with uncertain predictions flagged for future review.

### Application to Variants of Uncertain Significance

To assess the clinical utility of MoVUS, the model was applied to discovery datasets comprising VUS from ADRD-related ClinVar submissions and collaborator-provided cohorts from UCSF and CARD. We used probability thresholds of 0.1 and 0.9 for discovery to assess only very high confidence predictions. A summary of variant counts and classifications is provided in **Table 2**.

**Table 2.** MoVUS performance on external VUS datasets.

|  | Total Variants (N) | % Classified | Pathogenic | Benign | Unclassified |
| --- | --- | --- | --- | --- | --- |
| dbNSFP | 3271 | 61.6 | 368 | 1647 | 1256 |
| CARD | 58 | 63.8 | 16 | 21 | 21 |
| UCSF | 5 | 40 | 2 | 0 | 3 |

### Within dbNSFP

Given that a large majority of variants in dbNSFP are VUS without classification from HGMD or ClinVar, we decided to extract these variants as an internal discovery set. To focus discovery efforts on ADRD variants, we filtered for variants with ClinVar traits matching the following terms: ‘alzheim’ (Alzheimer’s), ‘dementia’ (generic dementia), ‘lewy’ (Lewy body dementia), ‘parkinson’ (Parkinson’s dementia), ‘frontotemp’, ‘ftd’, ‘Frontotemporal_dementia’ (FTD), ‘pick’ (Pick’s disease), ‘mixed’ (mixed dementia), and ‘creutzfeldt’, ‘jakob’ (CJD). This filtering resulted in 3,271 VUS variants. In total, 368 were classified as pathogenic, 1647 as benign, and 1256 remained unclassified.

### Collaborator Curated Variants

In the CARD variants set (n = 58), MoVUS classified 16 variants as pathogenic, 21 as benign, and 21 as uncertain. Of these 16 variants classified as pathogenic, 13 were excluded from further investigation as they were already classified as DM in HGMD. The two remaining variants, chr12:24842107:A:T near *BCAT1* and chr5:139307759:T:G near *MATR3*, were initially classified as “DM?” in HGMD and were newly classified as pathogenic by our model with predicted probabilities of 94% and 95%, respectively. Interestingly, one variant, chr6:146544084:C:G near *RAB32*, was previously classified as “DM” in HGMD, and our model predicted this variant as benign with a predicted probability of 100%.

In the UCSF VUS set (n = 5), two were classified as pathogenic, zero as benign, and three as uncertain. The variants classified as pathogenic were chr12:64474273:A:C and chr12:64466919:T:C in *TBK1* with predicted probabilities of 100% and 91%, respectively.

These results demonstrate the model’s ability to provide high probability predictions for a substantial proportion of VUS, while conservatively flagging uncertain cases.

### Clinical Utility

MoVUS provided high-confidence predictions (probability thresholds of 0.1 and 0.9) for an average of 55% of discovery variants across dbNSFP and collaborator-provided VUS, supporting its potential utility in clinical variant interpretation. SHAP-based local explanations were generated for individual VUS predictions, enabling assessment of the features driving each classification through our Streamlit web interface.

## Discussion

MoVUS outperforms any of the individual annotations on their own. While individual tools achieve a range of 56-94.5% balanced accuracy, with the highest being ClinPred scores, MoVUS’s ensemble approach provides more robust predictions across diverse variant types and clinical contexts. The model’s performance on independent test sets (91.5% recall for HGMD-only, 98.5% balanced accuracy for ClinVar-only) demonstrates improved generalizability compared to single-tool approaches. By creating a meta-consensus score that combines the best available evidence from multiple sources, MoVUS achieves higher accuracy than any single annotation while maintaining interpretability through SHAP analysis.

Importantly, MoVUS also demonstrated an ability to classify a substantial portion, 62%, of the 3,271 ADRD VUS from dbNSFP and >40% of collaborator curated VUS, providing new insights into previously uncharacterized genetic variation in neurodegenerative disease. We can move beyond benchmarking by applying MoVUS across collaborator discovery cohorts to evaluate real-world utility in functionally and clinically relevant datasets. For example, the pathogenic classification of *BCAT1* variant rs371256913 is supported by emerging evidence linking BCAT1 dysfunction to Alzheimer’s disease through autophagy regulation. Given that autophagy dysfunction is increasingly recognized as central to Alzheimer’s disease pathogenesis, the BCAT1 pathway represents a convergence point between metabolic dysfunction and protein aggregation, which supports the pathogenic classification of *BCAT1* variants in Alzheimer’s disease ^18^.

The RAB32 variant, chr6:146544084:C:G (p.Ser71Arg), presents an interesting case where MoVUS’s prediction (100% confidence benign) conflicts with its HGMD classification as DM. This variant has been associated with both familial and sporadic Parkinson’s disease, though it is reported with reduced penetrance ^19^. RAB32’s role in regulating LRRK2-mediated late endosomal trafficking relates to Parkinson’s disease pathogenesis ^20,21^. Interestingly, a recent study in a Chinese cohort found no association between RAB32 variants and Parkinson’s disease ^22^, and collaborators at UCSF found only one potential carrier of this variant within their genetic database at the MAC, who was a clinically normal, recruited control in their 80s. Inspection of another DM classified variant from HGMD with reduced penetrance, LRRK2 G2019S (chr12:40340400:G:A), similarly, is not recognized as pathogenic by MoVUS (predicted probability 0.547). The disparate classifications between HGMD and MoVUS here highlight that for variants with complex inheritance patterns due to reduced penetrance or population-specific factors, ensemble-based prediction algorithms may not capture variant deleteriousness. Nevertheless, MoVUS offers researchers an alternative classification framework that can inform a more nuanced interpretation of variant pathogenicity.

The pathogenic classified variant chr5:139307759:T:G in *MATR3* (p.Phe115Cys) was identified via exome sequencing in a European family with ALS and dementia and segregated with disease ^23^. Clinically, this variant is associated with a rapidly progressive phenotype, with affected individuals succumbing to respiratory failure within five years of onset ^23^. These findings support the pathogenic classification determined by MoVUS and demonstrate the clinical relevance of this variant in ALS pathogenesis.

The variants classified as pathogenic by MoVUS in *TBK1* were present in two patients at the UCSF MAC with confirmed clinical diagnoses of frontotemporal dementia. The patient with chr12:64474273:A:C was a Hispanic woman who presented with executive dysfunction, hoarding behavior, and episodic memory deficit, followed by asymmetric weakness and behavioral changes starting at age 63 years. Her clinical presentation could be explained by frontotemporal degeneration with motor neuron involvement consistent with prior reported cases with confirmed pathologic *TBK1* variants leading to TDP-43 proteinopathy. The carrier of chr12:64466919:T:C was a White woman who presented with gradually progressive symptoms consistent with semantic variant primary progressive aphasia associated with prominent behavioral deficits starting at age 60. Her autopsy at age 70 showed TDP 43, Type B neuropathology, consistent with prior pathological reports of pathogenic *TBK1* variant neuropathology (more details reported in a case series, with publication in process, of TBK1 mutation frontotemporal degeneration). These clinical features observed in the *TBK1* variant carriers support MoVUS’s pathogenic classification and the model’s ability to identify variants with well-characterized clinical and pathological correlates.

This study has several limitations. First, the training and evaluation datasets may lack sufficient ancestral and demographic diversity, potentially limiting the generalizability of MoVUS predictions to diverse populations. Second, the model is restricted to missense variants and does not address other variant types, such as noncoding or structural variants. This limitation reflects both the current lack of reference data from long-read sequencing and the fact that most annotation scores were developed specifically for missense variant classification. Third, MoVUS relies exclusively on sequence-based and functional prediction annotations; other potentially informative data types, such as epigenetic marks, regulatory annotations, or transcriptomic context, were not incorporated. Additionally, the model is agnostic to sex and does not account for potential sex-specific or ancestry-specific effects on variant pathogenicity.

A key limitation of our approach is the potential for circularity between training data and the development of individual functional prediction tools. Many tools included in MoVUS (such as REVEL, CADD, and MetaSVM) were trained on variants from HGMD and ClinVar, creating potential data leakage when these databases are used for model evaluation. To mitigate this concern, we implemented strict data splitting protocols and validated performance on truly independent collaborator datasets. At its core, MoVUS reduces potential bias by surveying all common annotation sources, an advantage over many publications that base concepts of variant pathogenicity on only one or a few.

Additionally, the high correlation between certain features (e.g., ClinPred and MutationTaster, r=0.87) raises concerns about double-counting similar predictive signals. While we removed features with correlation >0.9, the remaining correlations suggest some redundancy persists. We implemented LASSO regularization as an alternative feature selection method to validate our feature selection approach. LASSO selected all 25 features in our current set, independently validating our correlation-based feature selection methodology. The 100% overlap between LASSO-selected and our chosen features suggests that our feature set captures distinct predictive signals without significant redundancy.

Future work should address these limitations by incorporating more diverse datasets, expanding to additional variant types, and integrating broader categories of genomic and functional annotations. Developing ancestry-specific models and incorporating tissue-specific expression data could further improve prediction accuracy. Additionally, integration with clinical outcome data and longitudinal studies would enable validation of predictions in real-world clinical settings. Despite these limitations, MoVUS represents a significant advancement in variant classification, providing a robust foundation for future improvements in clinical variant interpretation.

In summary, MoVUS illustrates that combining diverse functional annotations through a balanced random forest approach can improve the classification of missense variants of uncertain significance. The model outperforms individual predictors and offers an interpretable framework that aids in triaging variants for follow-up in Alzheimer’s and related disorders. In the future, the MoVUS framework can be readily extended to incorporate new annotations to improve its predictions. By lowering barriers to variant interpretation, MoVUS helps in highlighting actionable candidates to accelerate discovery in neurodegenerative diseases.

## Acknowledgments

This research was supported by the Aligning Science Across Parkinson’s Initiative, the Intramural Research Program, National Institute on Aging, National Institutes of Health, Department of Health and Human Services, project ZO1 AG000949, and the Michael J. Fox Foundation for Parkinson’s Research. This work utilized the computational resources of the NIH STRIDES Initiative (https://cloud.nih.gov) through the Other Transaction agreement - Azure: OT2OD032100, Google Cloud Platform: OT2OD027060, Amazon Web Services: OT2OD027852. This work utilized the computational resources of the NIH HPC Biowulf cluster (https://hpc.nih.gov).

## Author contributions

Conceptualization: C.J., M.B.M., J.S.Y., A.B.S., M.A.N., H.L.L. Methodology: C.J., M.B.M., M.A.N., H.L.L. Investigation: C.J., M.B.M Visualization: C.J., M.B.M., Supervision: C.J., M.B.M., J.S.Y., A.B.S., M.A.N., H.L.L. Writing—original draft: C.J., M.B.M., J.S.Y., A.B.S., M.A.N., H.L.L. Writing—review & editing: C.J., M.B.M., J.S.Y., A.B.S., M.A.N., H.L.L., M.J.K.

## Supplementary information

If needed below.

